# idTAG enables programmable and physiologically faithful control of endogenous protein degradation

**DOI:** 10.64898/2026.09.18.752548

**Authors:** Mingkui San, Jiakun Peng, Fang Zhao, Rui Xiao

**Affiliations:** Department of Hematology, Medical Research Institute, Frontier Science Center for Immunology and Metabolism, Zhongnan Hospital of Wuhan University, Wuhan University, Wuhan 430071, Hubei, China; Department of Cardiology, Zhongnan Hospital of Wuhan University, Wuhan University, Wuhan 430071, Hubei, China

**Keywords:** idTAG, Tet-On regulation, Programmable protein degradation, RNA-binding proteins, Transcription regulation

## Abstract

Rapid and precise depletion of endogenous proteins is essential for dissecting gene function, yet existing degron technologies often require direct modification of endogenous coding sequences, which can compromise protein expression and limit their applicability. Here, we develop idTAG, an inducible degron platform that decouples protein expression from protein degradation by integrating Tet-On 3G-mediated transcriptional control with dTAG-mediated degradation. Systematic characterization reveals that conventional degron knock-in frequently causes unintended reduction of target protein expression at the mRNA level, representing an underappreciated limitation of current approaches. In contrast, idTAG enables efficient generation of degron cell lines while maintaining physiological protein abundance and allows programmable control of degradation kinetics through independent regulation of protein synthesis and degradation. Applying idTAG to RNA-binding proteins, we demonstrate that acute depletion of FUS has minimal effects on global RNA polymerase II transcription, whereas acute loss of SRSF5 selectively disrupts transcription termination at a subset of genes. Thus, idTAG provides a versatile platform for physiological and temporally controlled interrogation of endogenous protein functions.

**Highlights:**

- Endogenous degron knock-in frequently reduces target mRNA and protein abundance
- idTAG preserves physiological protein levels, enabling efficient generation of degron cell lines
- Independent control of mRNA expression and protein degradation of idTAG tunes depletion kinetics
- FUS loss has little effect on transcription, whereas SRSF5 loss selectively impairs transcription termination

**Graphical abstract:** 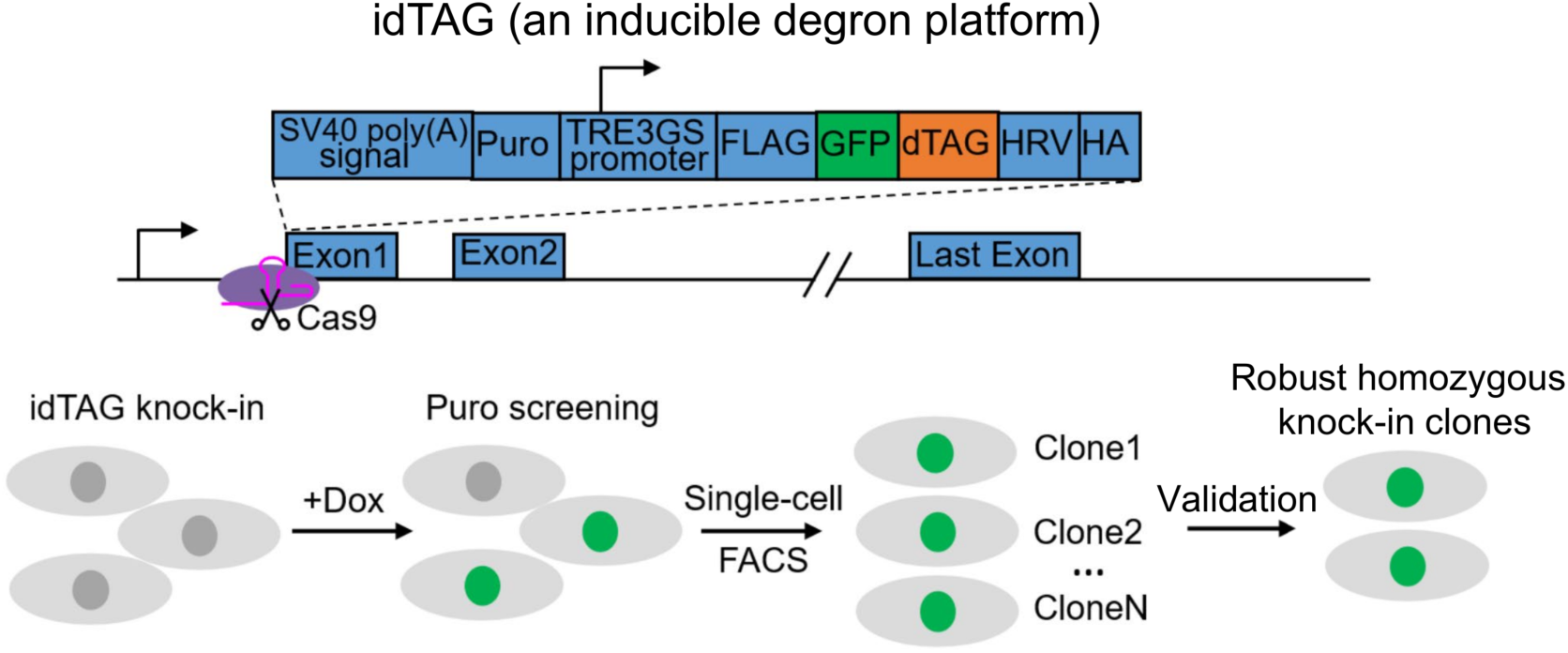

## Introduction

Precise manipulation of endogenous protein abundance is essential for dissecting gene function and establishing causal relationships between proteins and cellular phenotypes. Conventional genetic approaches, including knockout and RNA interference, often lack temporal control and may trigger compensatory responses that obscure direct molecular functions (El-Brolosy et al., 2019; El-Brolosy & Stainier, 2017). Recent advances in targeted protein degradation technologies, including auxin-inducible degron 2 (AID2), dTAG, and SD40 systems, have enabled rapid and selective depletion of endogenous proteins within minutes by harnessing ligand-induced ubiquitination and proteasomal degradation (Mercer et al., 2024; Nabet et al., 2018; Yesbolatova et al., 2020). These approaches have transformed the ability to interrogate protein function with unprecedented temporal resolution (Liang et al., 2023; Xin et al., 2026).

Despite their broad utility, current degron technologies commonly rely on direct insertion of degron tags into endogenous protein-coding sequences (Mercer et al., 2024; Nabet et al., 2018; Nishimura et al., 2009), which may introduce unintended consequences beyond ligand-induced degradation. In our systematic efforts to generate degron cell lines for RNA-binding proteins (RBPs), we found that endogenous degron knock-in frequently resulted in substantial reductions in target protein abundance and limited the recovery of homozygous degron clones. Consistent with our observations, recent studies have reported markedly reduced levels of endogenous degron-tagged proteins in mammalian systems, including CDK2-dTAG and CDK5-dTAG knock-in mice, as well as DNMT3B-AID2, U2AF2-AID2, and RBBP4-AID2 human iPSCs (Bondeson et al., 2022; Xing et al., 2025; Yenerall et al., 2023). Together, these findings reveal an underappreciated limitation of current degron strategies and highlight the need for approaches that enable controlled protein depletion while preserving physiological protein expression.

Overcoming this limitation is particularly important for studying RBPs, whose functions are frequently dynamic and context dependent. FUS, a member of the FET family of RBPs, is implicated in amyotrophic lateral sclerosis (ALS) and frontotemporal dementia (FTD), with disease-associated mutations causing nuclear depletion and cytoplasmic aggregation (Deng et al., 2014; Kwiatkowski et al., 2009; Vance et al., 2009). Previous studies using chronic FUS depletion have suggested that FUS regulates RNA polymerase II (Pol II)-dependent transcription, including control of Pol II CTD Ser2 phosphorylation (Schwartz et al., 2012). However, whether these effects represent direct consequences of acute FUS loss remains unclear.

Similarly, SRSF5, a member of the SR protein family best known for its role in pre-mRNA splicing (Li et al., 2023; Long & Caceres, 2008), may have additional transcriptional functions. Other SR proteins, including SRSF1 and SRSF2, directly regulate transcription through modulation of promoter-proximal pause release (Ji et al., 2013), whereas emerging evidence linking SRSF5 to the cleavage and polyadenylation machinery (Okuda et al., 2025) suggests a potential role in transcription termination. Nevertheless, the acute functions of SRSF5 in transcription remain unexplored.

Here, we develop an inducible degron platform, termed idTAG, that decouples protein expression from protein degradation by integrating Tet-On 3G-mediated transcriptional control with dTAG-mediated degradation. Unlike conventional degron knock-in strategies, idTAG enables efficient generation of degron cell lines while maintaining physiological protein abundance and provides programmable control of protein depletion kinetics. Applying idTAG to endogenous RBPs, we demonstrate that acute FUS depletion has minimal effects on global Pol II transcription, whereas acute SRSF5 depletion selectively disrupts transcription termination at a subset of genes. Thus, idTAG provides a general framework for physiological and temporally controlled interrogation of endogenous protein functions.

## Results

### Systematic generation of AID2 degron cell lines for RNA-binding proteins reveals a low success rate

To systematically investigate the transcriptional functions of chromatin-associated RNA-binding proteins (RBPs) (Xiao et al., 2019), we sought to establish AID2-based rapid degradation cell lines targeting 18 candidate RBPs in HepG2 cells. We first generated a parental HepG2 cell line stably expressing the engineered auxin receptor OsTIR1(F74G)-mCherry integrated at the AAVS1 safe-harbour locus (Figure 1A and B). Subsequently, for each candidate RBP, an mAID2–mEGFP–FLAG tagging cassette with a P2A-linked blasticidin-resistance marker was introduced at either the N or C terminus of the endogenous locus through CRISPR/Cas9-mediated homology-directed repair (HDR) (Qiu et al., 2022) (Figure 1A).

**Figure 1.**
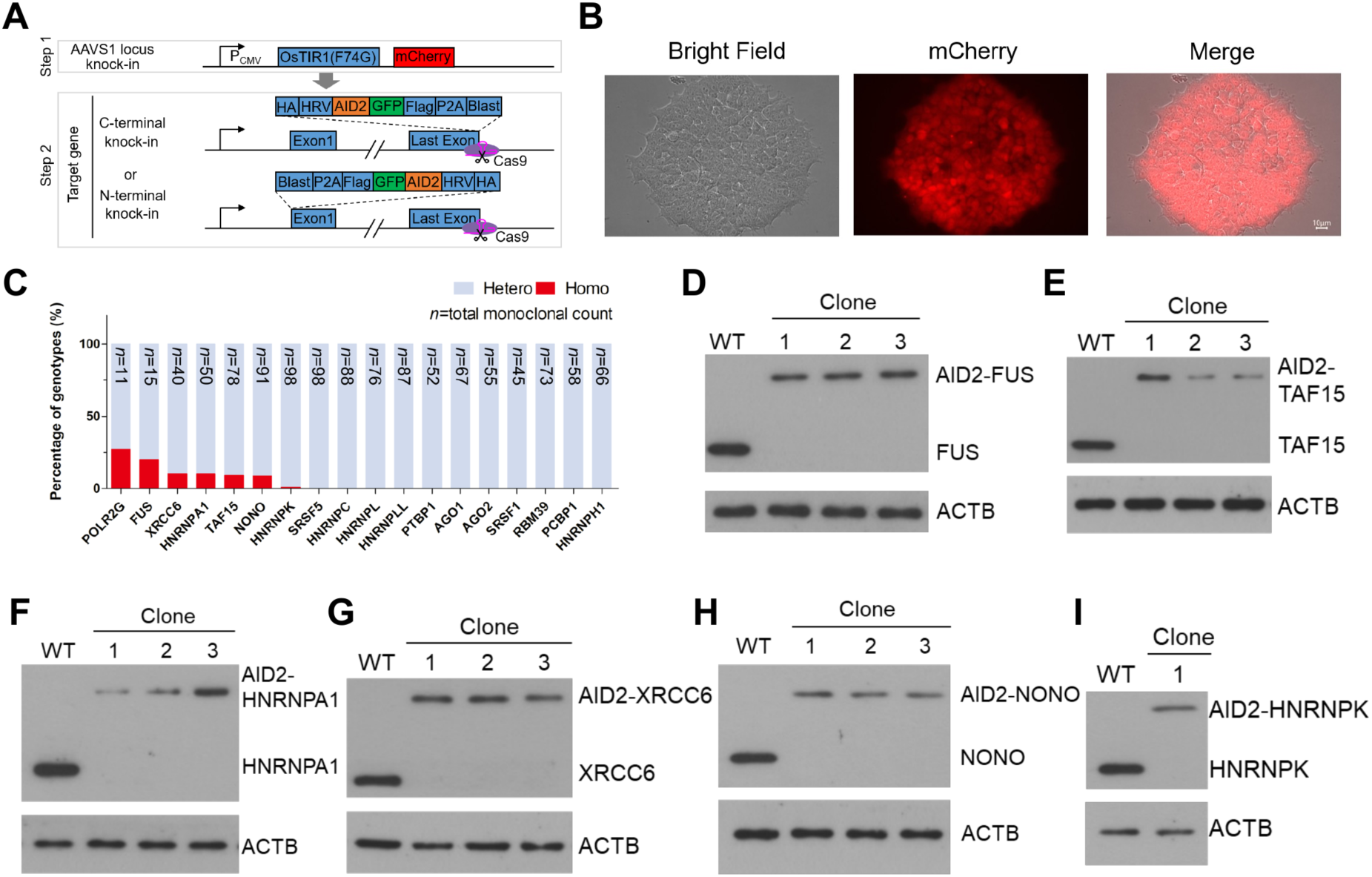
Systematic AID2 engineering of endogenous RNA-binding proteins reveals limited recovery of homozygous clones. A, Two-step strategy for generating endogenous AID2 cell lines. An OsTIR1(F74G)– mCherry expression cassette was first integrated at the AAVS1 safe-harbour locus, followed by CRISPR– Cas9-mediated insertion of an mAID2–mEGFP–FLAG tagging cassette with a P2A-linked blasticidin-resistance marker at the amino or carboxy terminus of the indicated target. B, Representative bright-field, mCherry and merged images of the OsTIR1(F74G)–mCherry parental line. Scale bar, 10 μm. C, Proportions of heterozygous and homozygous knock-in clones recovered for the 18 indicated RNA-binding proteins. The number above each bar is the total number of monoclonal lines screened for that target. D–I, Immunoblot validation of homozygous AID2 knock-in clones for FUS (D), TAF15 (E), HNRNPA1 (F), XRCC6 (G), NONO (H) and HNRNPK (I). Wild-type cells were analysed in parallel, and ACTB served as the loading control.

Following antibiotic selection, single-cell cloning, genotyping, and immunoblotting validation, we successfully obtained homozygous AID2 knock-in clones for only 7 of the 18 targeted RBPs, including POLR2G, FUS, XRCC6, HNRNPA1, TAF15, NONO, and HNRNPK (Figure 1C–1I and <u>Figure S1A–F</u>). In contrast, homozygous knock-in clones could not be generated for the remaining 11 targets, including SRSF5, HNRNPC, and AGO2, despite extensive screening. For example, even after isolating 98 independent single-cell clones for SRSF5, no homozygous knock-in clone was obtained (Figure 1C).

**Figure S1.**
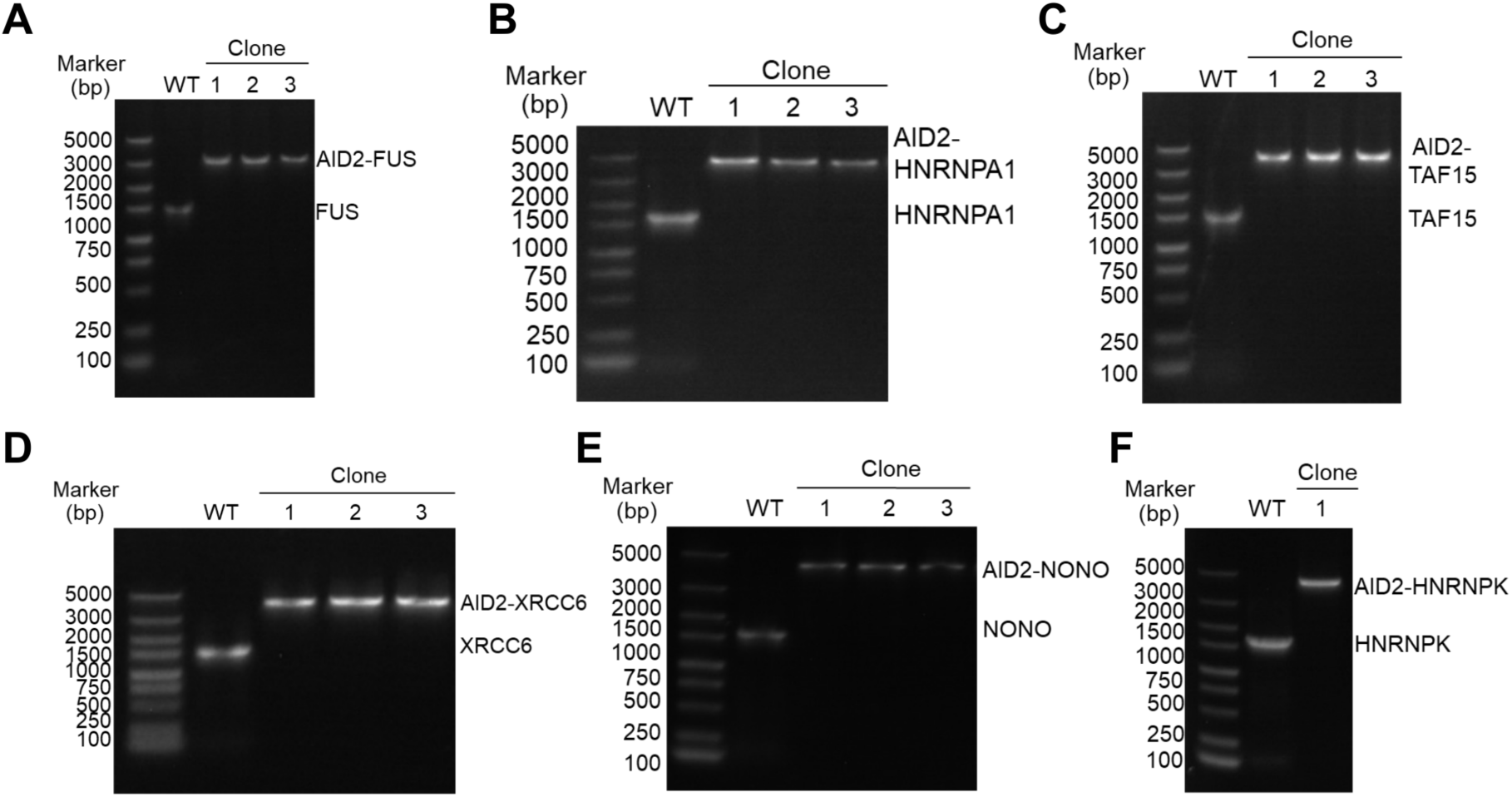
Genomic PCR validation of homozygous AID2 knock-in clones. A–F, Genomic PCR analysis of wild-type cells and candidate homozygous AID2 knock-in clones for *FUS* (A), *HNRNPA1* (B), *TAF15* (C), *XRCC6* (D), *NONO* (E) and *HNRNPK* (F). Amplicons corresponding to the tagged and unmodified alleles are indicated; marker sizes are shown in base pairs.

Notably, even among successfully established homozygous knock-in lines, the abundance of AID2-tagged proteins was substantially lower than that of the corresponding endogenous proteins in wild-type (WT) cells. For instance, FUS protein levels were consistently and significantly lower in three independent homozygous clones carrying the C-terminal AID2 cassette than in WT cells (Figure 1D). Similar reductions in protein abundance were observed for HNRNPA1, TAF15, NONO, and HNRNPK (Figure 1E, 1F, 1H, and 1I), consistent with previous observations that degron tagging can affect basal protein stability or expression levels (Bondeson et al., 2022; Xing et al., 2025; Yenerall et al., 2023).

Together, these findings indicate that endogenous AID2 tagging can impose an unintended basal destabilization effect on target proteins, limiting the generation of degron cell lines for a substantial fraction of RBPs. This effect may be particularly detrimental for essential genes, where even moderate reductions in protein abundance could compromise cellular fitness and prevent the recovery of viable homozygous degron clones (Hart et al., 2015; Li et al., 2024) (Figure 1C). These observations highlight an important limitation of current degron strategies and underscore the need for improved approaches that enable high-efficiency generation of degron cell lines while preserving physiological protein expression and function.

### Endogenous degron tagging unexpectedly reduces target gene expression

To investigate the mechanism underlying reduced protein abundance following degron tagging, we first examined whether the position of degron fusion influenced target protein expression using FUS as a model. We generated a cell line carrying an N-terminal AID2 fusion of endogenous FUS (Figure 2A). Although C-terminal AID2 fusion caused a more pronounced reduction in FUS protein abundance than did N-terminal fusion (Figure 2B and C), both N- and C-terminal AID2 tagging substantially decreased FUS protein levels. These results indicate that degron fusion at either terminus can adversely affect endogenous protein expression, with C-terminal tagging exerting a stronger effect.

**Figure 2.**
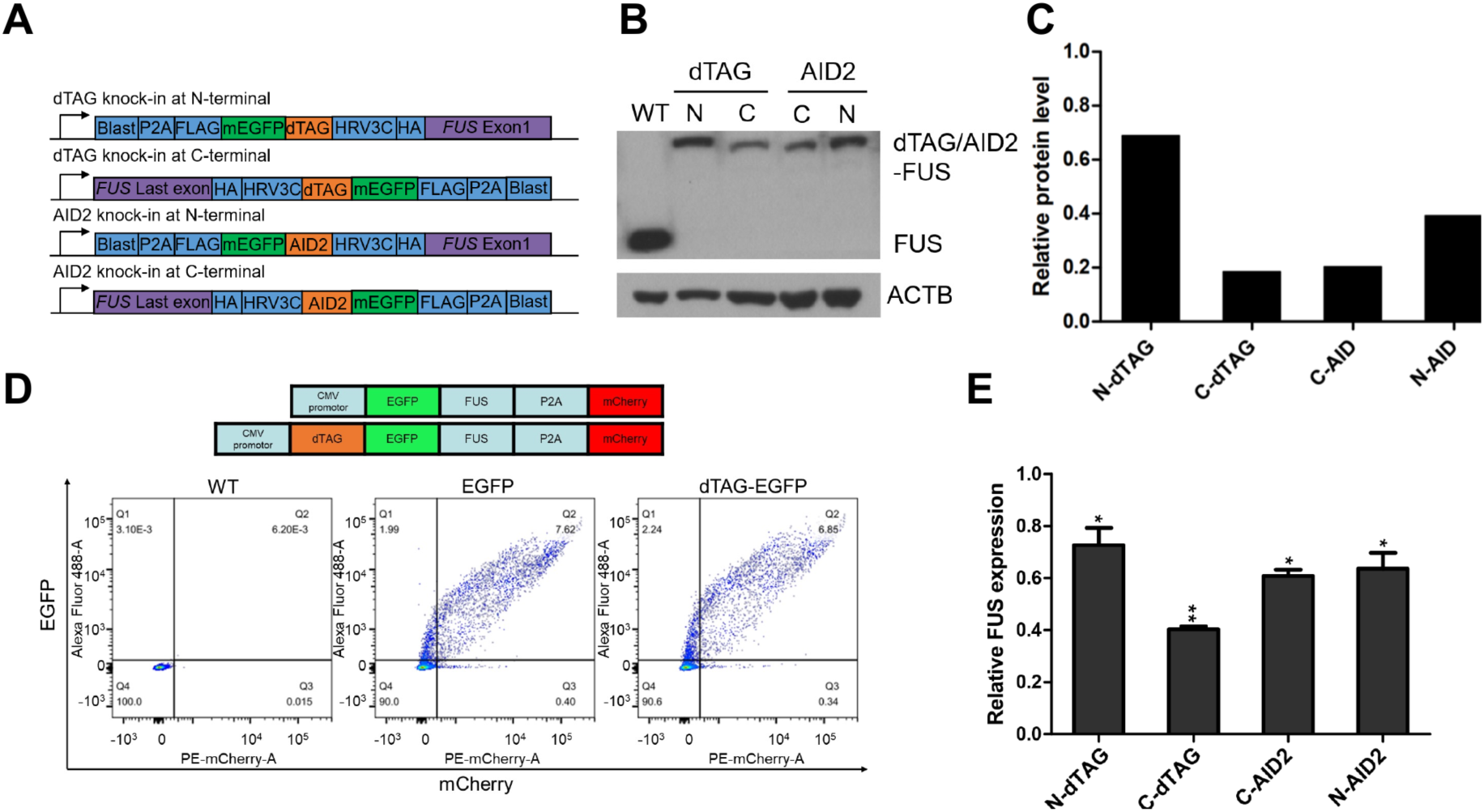
Endogenous degron knock-in reduces FUS expression independently of basal dTAG-mediated degradation. A, Schematics of amino- and carboxy-terminal dTAG or AID2 knock-in configurations at the endogenous *FUS* locus. B, Immunoblot analysis of wild-type cells and the four indicated homozygous knock-in lines. Tagged FUS migrates above unmodified FUS; ACTB is the loading control. C, Densitometric quantification of the immunoblot in B after normalization to ACTB and to wild-type FUS. D, P2A-based dual-fluorescence reporter design and representative flow-cytometry plots. EGFP reports the abundance of EGFP–FUS or FKBP12(F36V)–EGFP–FUS, whereas mCherry provides a co-translated internal control. E, RT–qPCR analysis of endogenous *FUS* transcript abundance in the indicated knock-in lines. Values were normalized to *ACTB* and to wild-type cells. Data are mean ± s.d. from three independent experiments. Statistical significance was assessed against wild type by an unpaired two-tailed Student’s t-test; *P < 0.05 and **P < 0.01.

We next asked whether this phenomenon was specific to AID2 or represented a general consequence of endogenous degron tagging. To this end, we generated FUS degron cell lines carrying dTAG fusions at either the N or C terminus (Figure 2A). Similar to AID2 tagging, both N- and C-terminal dTAG fusions resulted in significant reductions in FUS protein abundance, with a stronger effect observed for C-terminal fusion (Figure 2B and C). Together, these results suggest that endogenous degron knock-in can broadly impair target protein expression, and that the extent of reduction is influenced by the position of tag insertion.

Previous studies have suggested that certain degron tags may induce basal degradation of target proteins even in the absence of the inducing ligand (Bondeson et al., 2022; Li et al., 2019; Yesbolatova et al., 2020). To determine whether intrinsic tag-mediated instability contributed to the reduced FUS abundance observed in our degron lines, we established a dual-fluorescence reporter system consisting of an EGFP-FUS-P2A-mCherry cassette, in which the EGFP/mCherry ratio serves as a proxy for FUS protein stability (Figure 2D). Quantification by flow cytometry revealed that dTAG fusion did not alter the EGFP/mCherry ratio in the absence of dTAG-13 treatment (Figure 2D), arguing against intrinsic dTAG-mediated destabilization as the primary cause of reduced FUS abundance.

We therefore examined whether degron knock-in affected *FUS* transcript levels. Unexpectedly, RT-qPCR analysis revealed a consistent 30–60% reduction in *FUS* mRNA abundance across all four degron cell lines, which closely correlated with the reduction in FUS protein levels (Figure 2C and E). These findings indicate that the reduced protein abundance caused by endogenous degron tagging primarily results from decreased target gene expression at the transcript level rather than ligand-independent protein degradation. We speculate that CRISPR/Cas9-mediated genome editing during degron insertion may induce locus-specific alterations in chromatin organization or three-dimensional genome architecture, thereby affecting endogenous gene expression (Bantele et al., 2025; Wang et al., 2025).

### Development of the idTAG inducible degradation system

To overcome the unintended reduction in endogenous protein expression caused by conventional degron knock-in strategies, we developed an inducible degron system, termed idTAG, which integrates Tet-On 3G-mediated transcriptional control with dTAG-mediated protein degradation (Figure 3A). In contrast to conventional approaches that fuse degron tags directly to the endogenous coding sequence, idTAG introduces an inducible expression cassette into the 5′ untranslated region (5′ UTR) of the target gene. This cassette consists of an SV40 polyadenylation signal to terminate transcription from the endogenous promoter, a puromycin resistance cassette for selection, a TRE3GS doxycycline (Dox)-responsive promoter, and a FLAG–mEGFP–FKBP12(F36V)–HRV–HA-tagged open reading frame (Figure 3A). Consequently, expression of the degron-tagged protein is placed under exclusive control of Dox induction.

**Figure 3.**
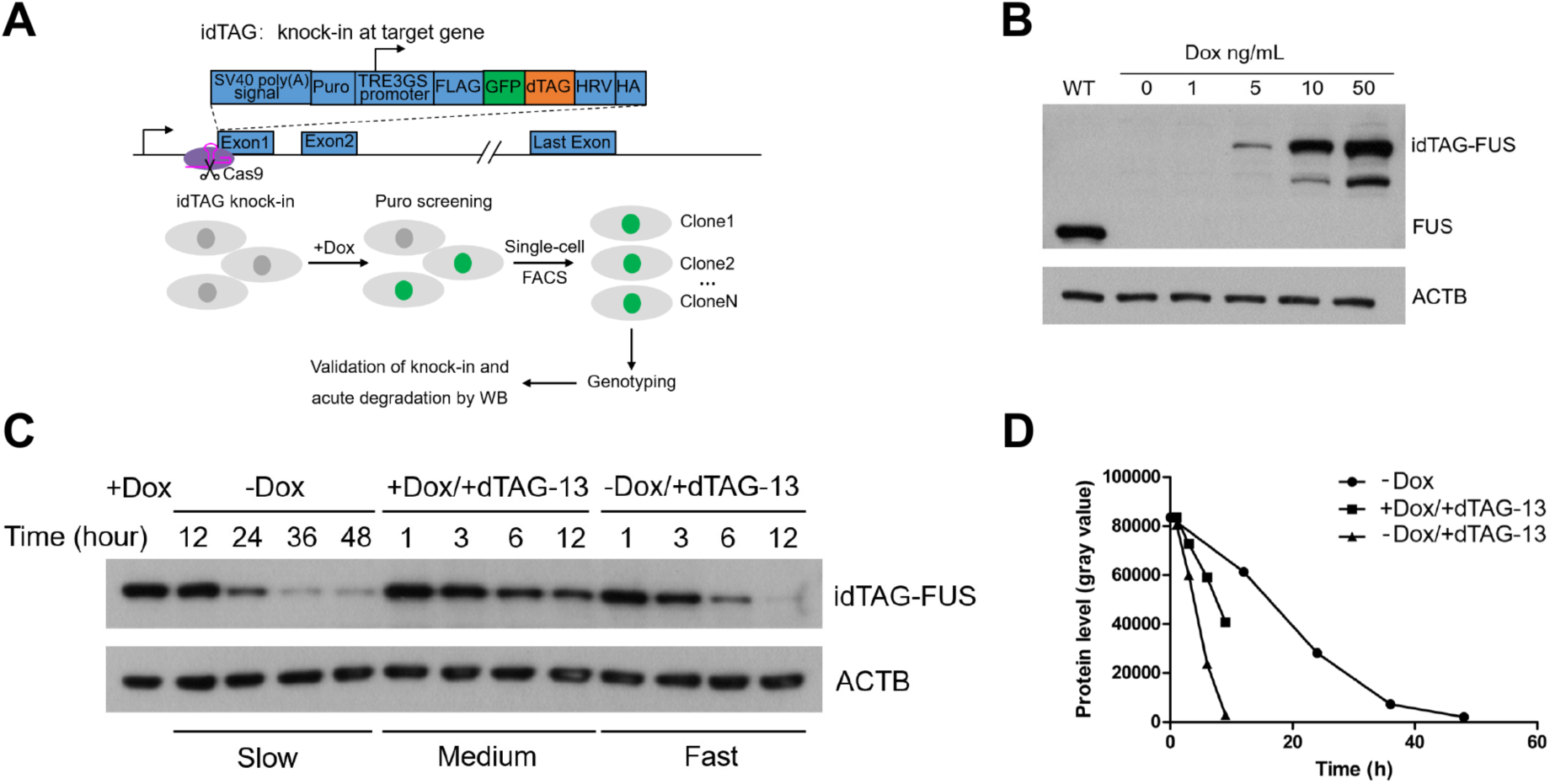
idTAG restores physiological FUS abundance and enables tunable degradation kinetics. A, Design and workflow of idTAG. A promoter-insulating SV40 poly(A) signal, puromycin-resistance cassette and TRE3GS promoter were inserted upstream of a FLAG-mEGFP-FKBP12(F36V)-HRV-HA-tagged target gene. Doxycycline (Dox) drives target-gene synthesis, and dTAG-13 promotes degradation of the FKBP12(F36V)-tagged protein. B, Immunoblot analysis of FUS-idTAG cells cultured for 48 h with the indicated Dox concentrations. Wild-type FUS and ACTB are shown as controls. C, Immunoblot analysis of three FUS-depletion modes after pre-equilibration with 10 ng/mL Dox for 48 h: slow depletion by Dox withdrawal alone; intermediate depletion by maintaining Dox and adding 0.5 μM dTAG-13; and rapid depletion by Dox withdrawal combined with 0.5 μM dTAG-13. Samples were collected at the indicated times, and ACTB served as the loading control. D, Densitometric time courses corresponding to C, based on ACTB-normalized FUS band intensities (arbitrary units).

To generate idTAG cell lines, cells undergoing targeted integration were selected with puromycin, followed by GFP-based fluorescence-activated cell sorting (FACS), single-cell cloning, genotyping, and immunoblot validation (Figure 3A). Using FUS as a model, genomic PCR confirmed homozygous insertion of the idTAG cassette (<u>Figure S2</u>). In the absence of Dox, FUS-idTAG cells exhibited undetectable expression of the tagged FUS protein, whereas Dox treatment restored FUS expression in a dose-dependent manner over a concentration range of 1–50 ng/mL. Importantly, treatment with 10 ng/mL Dox restored FUS protein abundance to a level comparable to that of endogenous FUS in wild-type cells (Figure 3B). These results demonstrate that idTAG enables precise restoration of physiological protein expression levels while avoiding the unintended consequences associated with direct degron fusion to endogenous proteins.

**Figure S2.**
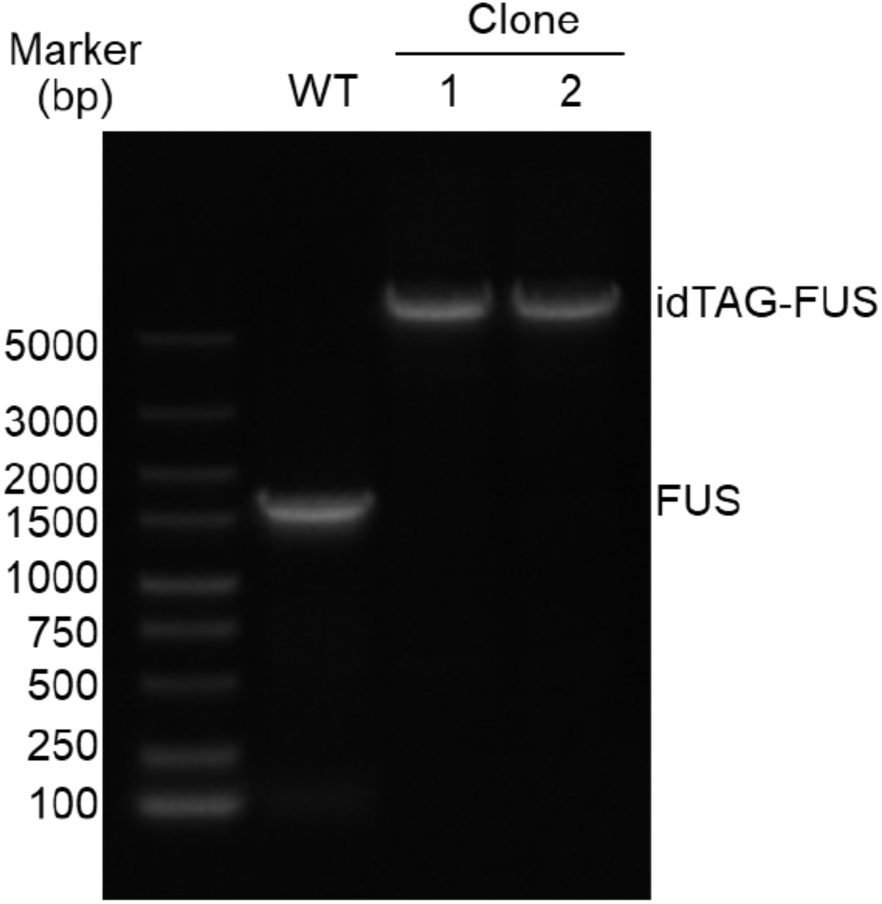
Genomic PCR validation of the FUS-idTAG knock-in cell line. Genomic PCR analysis of wild-type cells and two homozygous FUS-idTAG clones. Amplicons corresponding to the tagged and unmodified alleles are indicated; marker sizes are shown in base pairs.

Beyond programmable expression control, idTAG also enables tunable regulation of protein degradation kinetics through independent modulation of Dox and dTAG-13. Using FUS-idTAG cells pre-induced with 10 ng/mL Dox, we established three distinct degradation modes (Figure 3C and D). Withdrawal of Dox alone resulted in slow depletion of FUS, with FUS abundance falling to approximately 50% at around 20 h and ∼20% remaining after 48 h (Figure 3C, left; Figure 3D). Maintaining Dox supplementation while adding 0.5 μM dTAG-13 induced moderate degradation, reducing FUS abundance to ∼50% within 12 hours (Figure 3C, middle; Figure 3D). In contrast, combining Dox withdrawal with dTAG-13 treatment triggered rapid depletion, achieving >80% FUS loss within 6 hours (Figure 3C, right; Figure 3D). Together, these results establish idTAG as a versatile platform that enables both physiological restoration and programmable degradation of endogenous proteins.

### Acute FUS depletion has minimal impact on global transcription

Several ALS-associated *FUS* mutations are linked to reduced nuclear FUS abundance and cytoplasmic aggregation, which are thought to contribute to disease pathogenesis (Deng et al., 2014; Kwiatkowski et al., 2009; Vance et al., 2009). In addition, FUS ChIP-seq analysis has revealed extensive association of FUS with promoter-proximal regions across the human genome (Xiao et al., 2019), while chronic FUS depletion by RNA interference has been reported to affect RNA polymerase II (Pol II)-mediated transcription (Schwartz et al., 2012). These observations have suggested that FUS-dependent transcriptional regulation may contribute to ALS pathology. However, whether acute loss of FUS directly affects transcription remains unclear.

To address this question, we used FUS-idTAG cells to examine the immediate consequences of FUS depletion. Cells were pre-equilibrated with 10 ng/mL Dox to restore physiological FUS expression, followed by Dox withdrawal combined with 0.5 μM dTAG-13 treatment for 6 hours to achieve rapid FUS depletion. We then performed Pol II ChIP-seq and global run-on sequencing (GRO-seq) analyses to assess transcriptional consequences following acute FUS loss. Biological replicates showed high concordance for both POLR2A ChIP–seq and GRO-seq (<u>Figure S3</u>).

**Figure S3.**
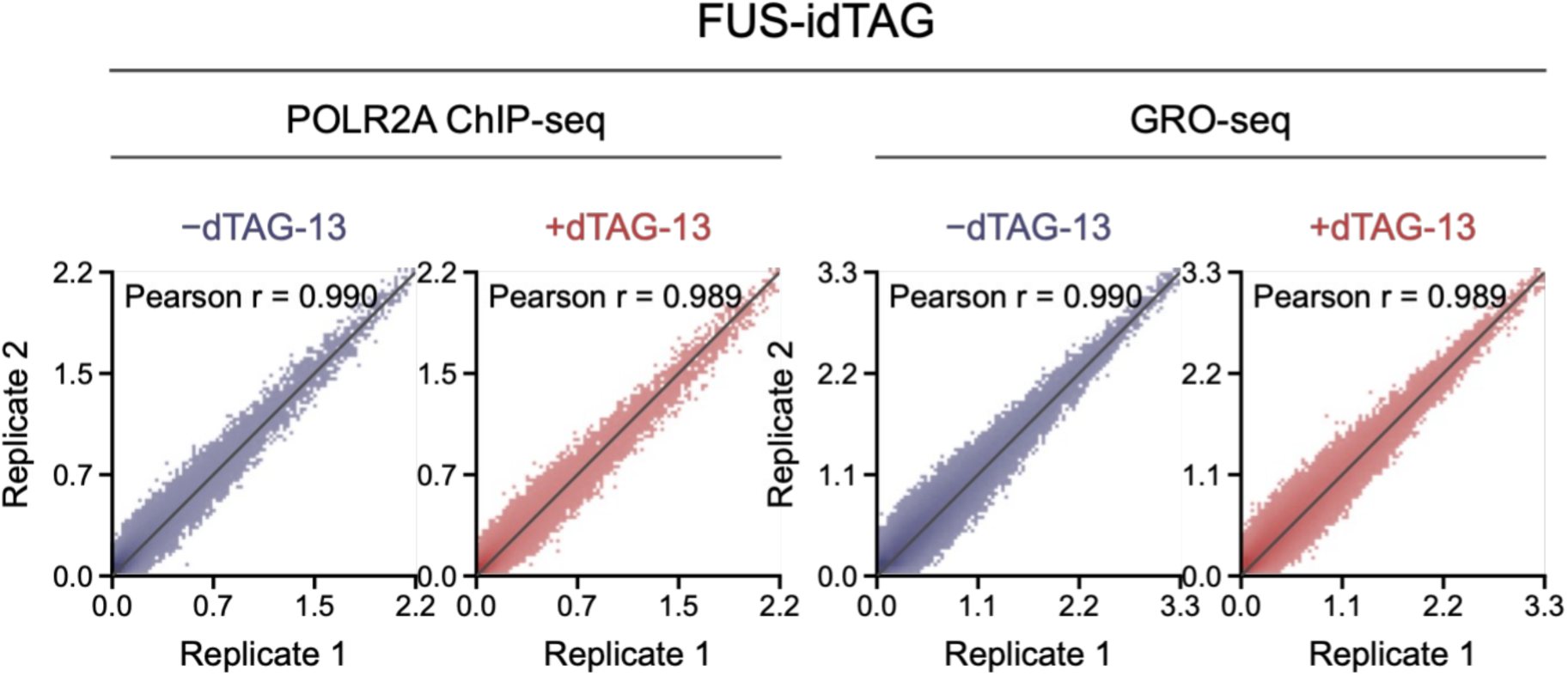
Biological-replicate concordance for genomic assays in FUS-idTAG cells. Replicate correlations for POLR2A ChIP-seq and GRO-seq in non-depleted and acutely FUS-depleted FUS-idTAG cells. Correlations were calculated from non-overlapping 1-kb genomic bins; Pearson correlation coefficients are shown. Each condition comprised two biological replicates.

Unexpectedly, acute FUS depletion had minimal effects on transcriptional activity. At representative loci, including *RPL13* and *MRPL36-NDUFS6*, Pol II ChIP-seq profiles remained largely unchanged following acute FUS degradation. Similarly, GRO-seq signals were unaffected at these loci (Figure 4A). Genome-wide metagene analysis further revealed only marginal changes in Pol II occupancy across transcribed regions upon acute FUS depletion (Figure 4B). Consistently, neither the pause release ratio (PRR; Kolmogorov–Smirnov test, *P* = 0.14) nor the readthrough index (RTI; Kolmogorov–Smirnov test, *P* = 0.77) showed appreciable changes (Figure 4C and D). GRO-seq analysis further confirmed that nascent transcription profiles remained largely unchanged genome-wide (Figure 4E–G).

**Figure 4.**
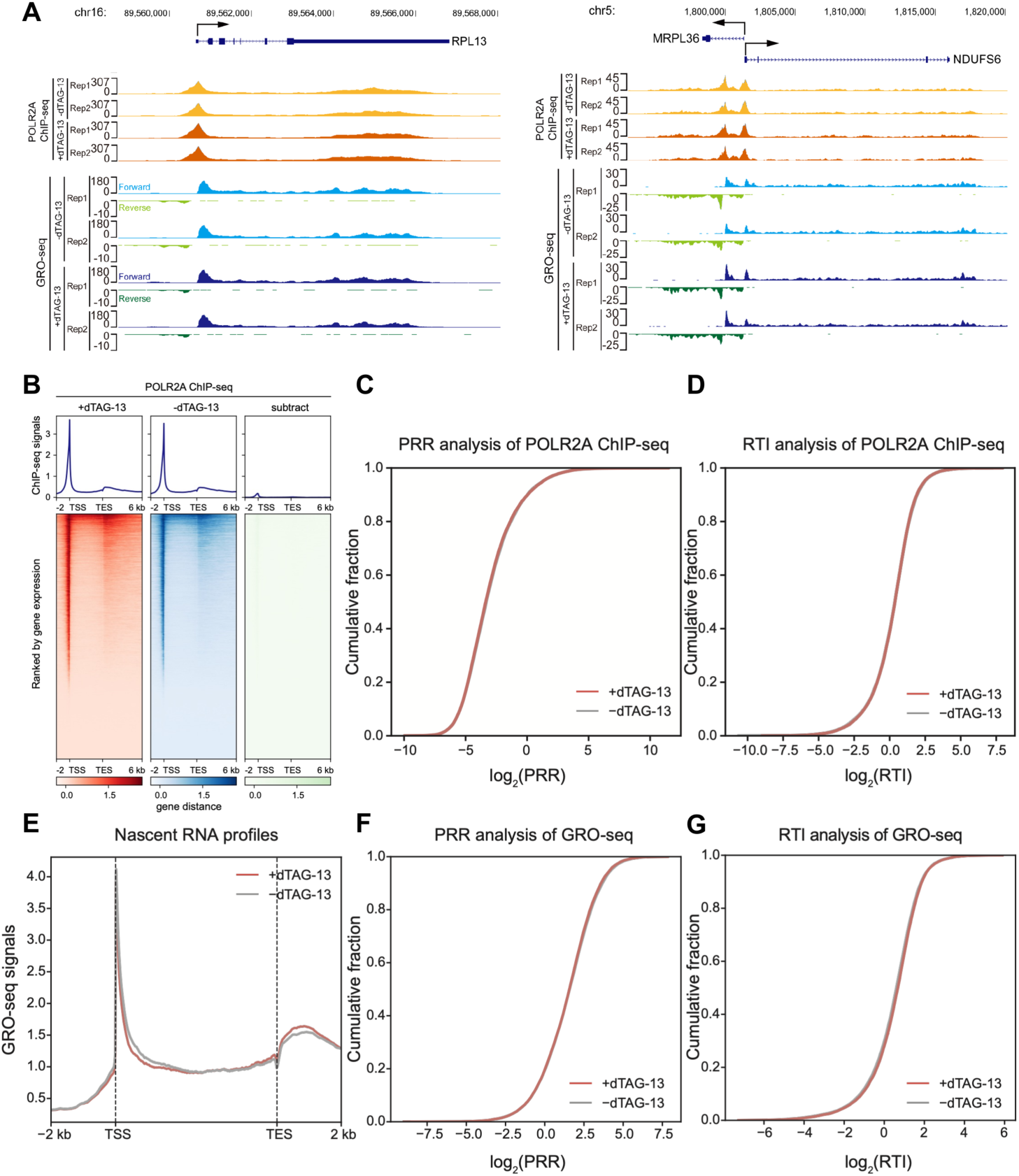
Acute FUS depletion causes little global change in POLR2A occupancy or nascent transcription. A, Genome-browser tracks for POLR2A ChIP–seq and strand-resolved GRO-seq at *RPL13* and the divergent *MRPL36*–*NDUFS6* locus before and after acute depletion. Two biological replicates are shown for each condition and assay. B, Average profiles and heat maps of POLR2A ChIP–seq signal from 2 kb upstream of the transcription start site (TSS) to 6 kb downstream of the transcription end site (TES) in depleted cells, control cells and the depleted-minus-control difference. C, Empirical cumulative distribution of log_2_ pause-release ratios (PRRs) from POLR2A ChIP–seq for 11,649 genes with control GRO-seq TPM > 1 (two-sample Kolmogorov–Smirnov test, D = 0.015, P = 0.14). D, Empirical cumulative distribution of the POLR2A ChIP–seq readthrough index (RTI) for 11,902 genes with control GRO-seq TPM > 1 (two-sample Kolmogorov–Smirnov test, D = 0.009, P = 0.77). E, Mean sense GRO-seq profiles across 14,788 protein-coding genes from 2 kb upstream of the TSS to 2 kb downstream of the TES. F, Empirical cumulative distribution of GRO-seq PRR for 11,887 genes with control TPM > 1 (two-sample Kolmogorov–Smirnov test, D = 0.015, P = 0.16). G, Empirical cumulative distribution of the GRO-seq RTI for 12,337 genes with control TPM > 1 (two-sample Kolmogorov–Smirnov test, D = 0.031, P = 1.9 × 10^−5^). NC, non-depleted control; KD, acute depletion. RTI calculations were adapted from a published RBM22 analysis (Du et al., 2024); signal from the TES to 2 kb downstream was normalized to gene-body signal for POLR2A ChIP–seq and to terminal-exon signal for GRO-seq.

Together, these results demonstrate that acute and near-complete depletion of FUS does not substantially perturb global Pol II transcription. These findings suggest that the transcriptional consequences of FUS loss may be context-dependent and that previously reported effects from chronic FUS depletion may involve secondary responses or long-term adaptation.

### SRSF5 preferentially associates with promoter-proximal chromatin regions

Although we were unable to generate SRSF5-AID2 knock-in cells using conventional degron engineering, we successfully established SRSF5-idTAG cells using the idTAG system (<u>Figure</u> <u>S4A</u>). Importantly, SRSF5-idTAG cells maintained SRSF5 protein expression at a level comparable to that in wild-type (WT) cells and enabled efficient acute depletion of SRSF5 within 3 hours following Dox withdrawal and dTAG-13 treatment (Figure 5A and B). These results demonstrate that idTAG enables functional interrogation of proteins that are otherwise difficult to study using conventional degron approaches.

**Figure 5.**
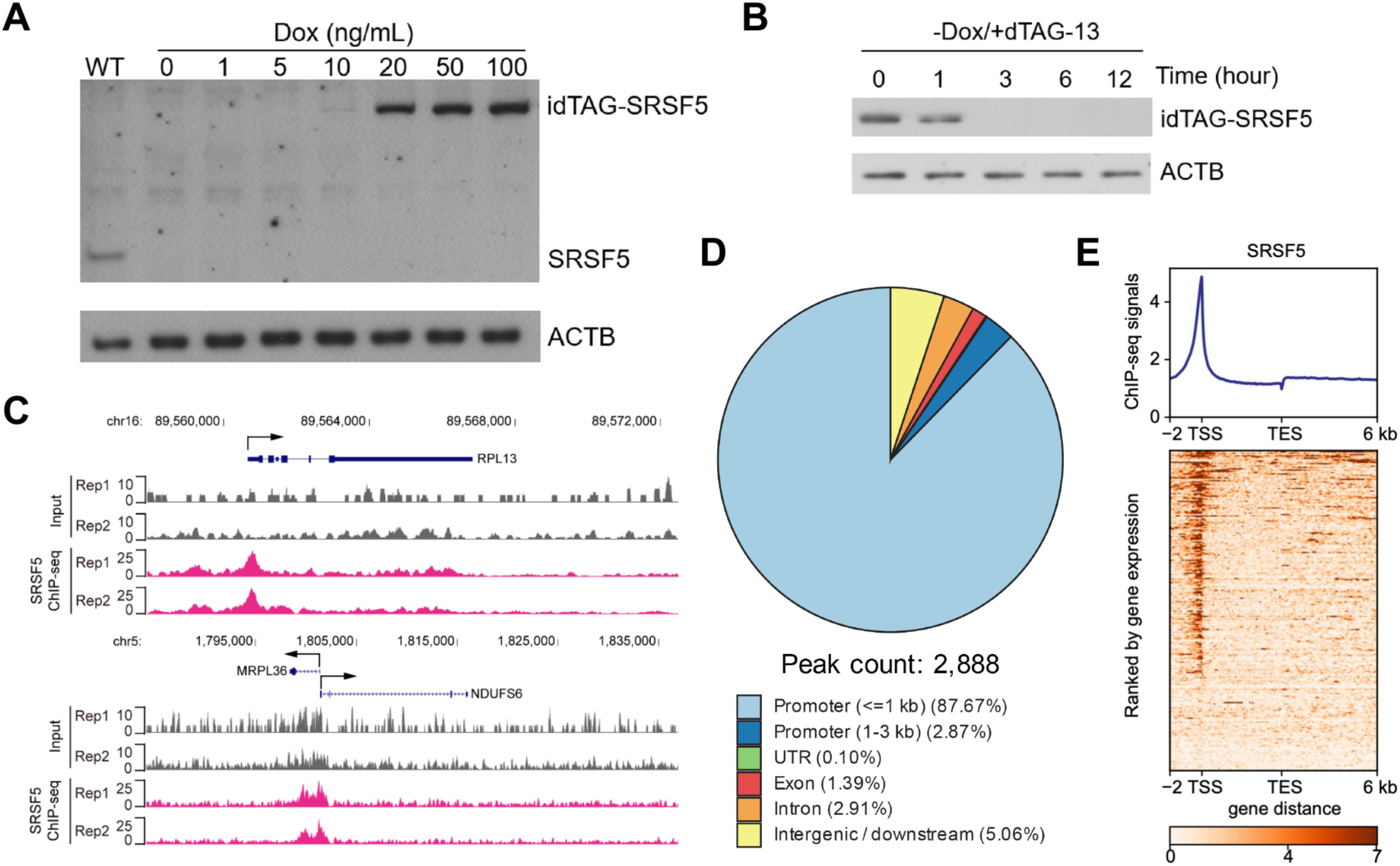
Generation of SRSF5-idTAG cells and genome-wide mapping of SRSF5 chromatin occupancy. A, Immunoblot analysis of SRSF5-idTAG cells cultured for 48 h with 0-100 ng/mL Dox. Wild-type SRSF5 and ACTB are shown as controls. B, Time course of rapid SRSF5 depletion after Dox withdrawal and addition of 0.5 μM dTAG-13. ACTB is the loading control. C, Input and FLAG ChIP-seq tracks from two biological replicates at *RPL13* and the divergent *MRPL36*-*NDUFS6* locus. D, Genomic annotation of 2,888 pooled SRSF5 peaks called at q < 0.05 after blacklist filtering. Peaks were assigned to promoters within 1 kb of a TSS (87.67%), promoters 1-3 kb from a TSS (2.87%), UTRs (0.10%), exons (1.39%), introns (2.91%) or intergenic/downstream regions (5.06%). E, Average profile and heat map of SRSF5 ChIP-seq signal across protein-coding genes from 2 kb upstream of the TSS to 6 kb downstream of the TES.

To investigate the chromatin association profile of SRSF5, we performed SRSF5 ChIP-seq analysis using SRSF5-idTAG cells, in which endogenous SRSF5 was fused with a FLAG epitope tag (Figure 3A). Biological replicates showed high concordance (<u>Figure S4B</u>). At representative loci, including *RPL13* and *NDUFS6*, SRSF5 exhibited strong enrichment at promoter-proximal regions (Figure 5C). Genome-wide analysis identified 2,888 high-confidence SRSF5-binding peaks, of which 87.67% were located within promoter regions (Figure 5D). Consistently, metagene analysis revealed pronounced SRSF5 enrichment around transcription start sites (TSSs), with stronger occupancy associated with actively transcribed genes (Figure 5E).

**Figure S4.**
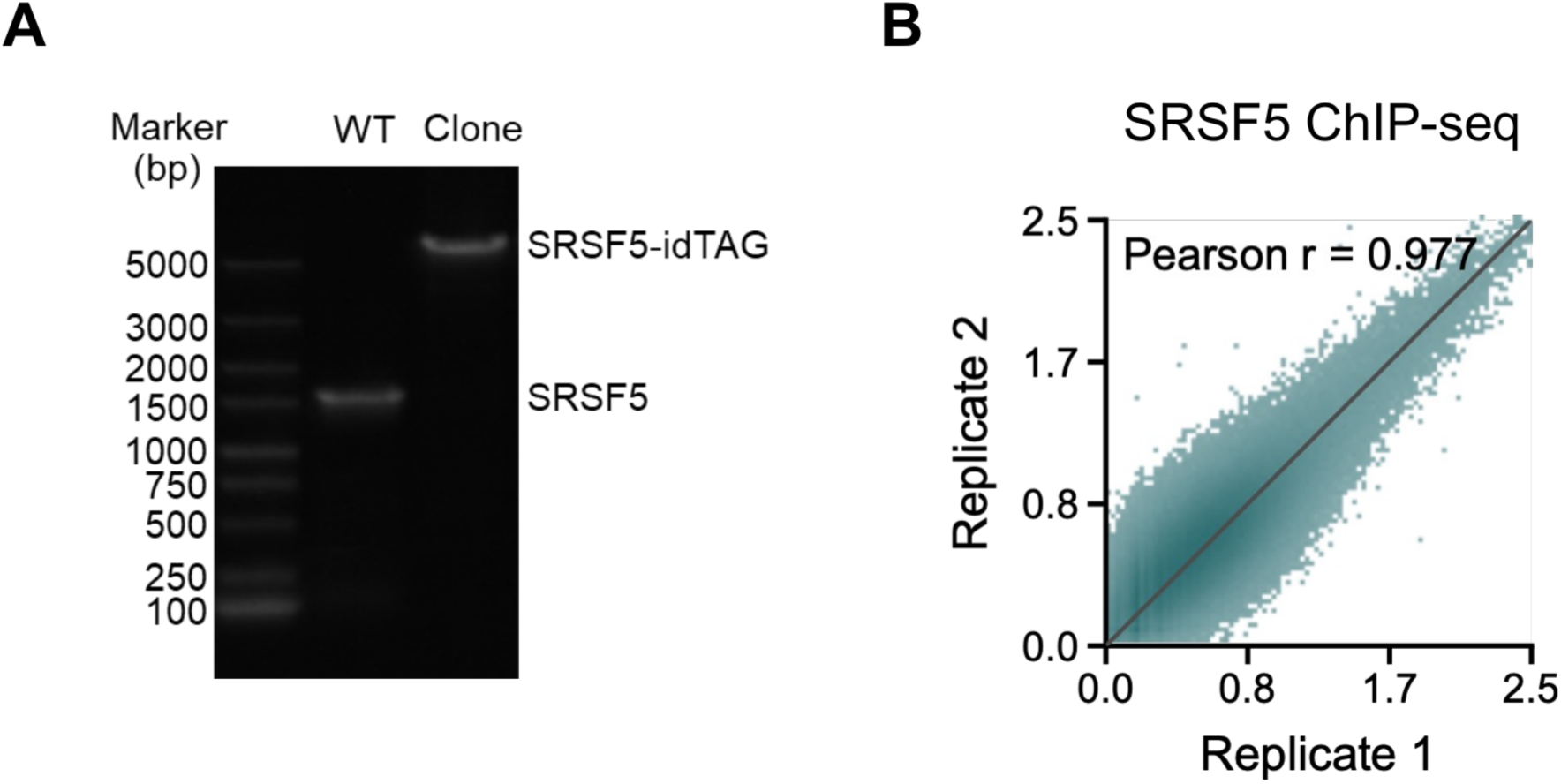
Genomic PCR validation and SRSF5 ChIP-seq replicate concordance in SRSF5-idTAG cells. A, Genomic PCR analysis of wild-type cells and a homozygous SRSF5-idTAG clone. Amplicons corresponding to the tagged and unmodified alleles are indicated; marker sizes are shown in base pairs. B, Correlation between two biological replicates of SRSF5 FLAG ChIP-seq, calculated from non-overlapping 1-kb genomic bins. The Pearson correlation coefficient is shown.

Similar preferential association of SR proteins with promoter-proximal chromatin regions has been reported for SRSF1 and SRSF2 (Ji et al., 2013), suggesting that promoter-proximal chromatin localization may represent a conserved feature of SR protein biology.

### SRSF5 selectively regulates transcription termination at a subset of genes

We next investigated the functional consequences of acute SRSF5 depletion using SRSF5-idTAG cells. Following acute SRSF5 degradation (Dox withdrawal combined with 0.5 μM dTAG-13 treatment for 6 hours), we performed POLR2A ChIP-seq to measure total RNA polymerase II (Pol II) occupancy, POLR2A-S2P ChIP-seq to monitor elongating Pol II, and GRO-seq to assess nascent transcription. Biological replicates showed high concordance under both control and SRSF5-depleted conditions (<u>Figure S5</u>).

**Figure S5.**
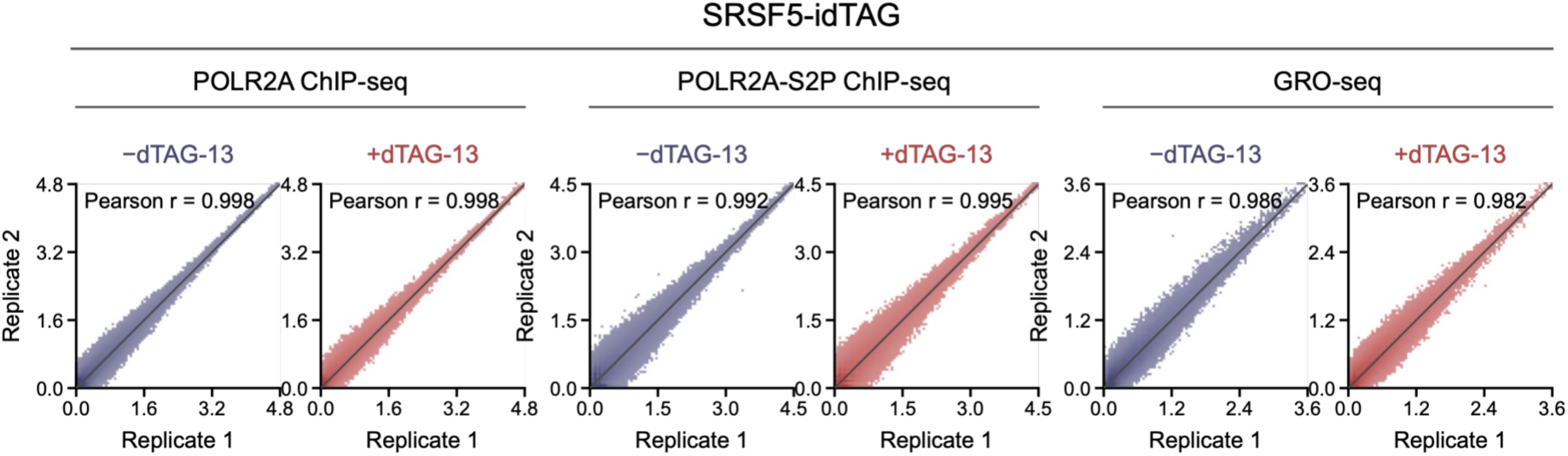
Biological-replicate concordance for genomic assays in SRSF5-idTAG cells. Replicate correlations for POLR2A ChIP-seq, POLR2A-S2P ChIP-seq and GRO-seq in non-depleted and acutely SRSF5-depleted SRSF5-idTAG cells. Correlations were calculated from non-overlapping 1-kb genomic bins; Pearson correlation coefficients are shown. Each condition comprised two biological replicates.

At representative loci, including *RPL13* and *MRPL36-NDUFS6*, total Pol II occupancy, elongating Pol II signals, and nascent transcription profiles remained largely unchanged following acute SRSF5 depletion (Figure 6A). Genome-wide metagene analysis further revealed only minimal changes in Pol II distribution across gene bodies upon SRSF5 loss (Figure 6B). Consistently, the pause release ratio (PRR) showed no appreciable global shift (two-sample Kolmogorov–Smirnov test, *P* = 1.51 × 10^−1^; Figure 6C), and GRO-seq analysis likewise revealed no appreciable global shift in transcriptional output or PRR (Figure 6D and 6E). Together, these results indicate that, unlike SRSF1 and SRSF2 (Ji et al., 2013), SRSF5 does not broadly affect transcription initiation or elongation.

**Figure 6.**
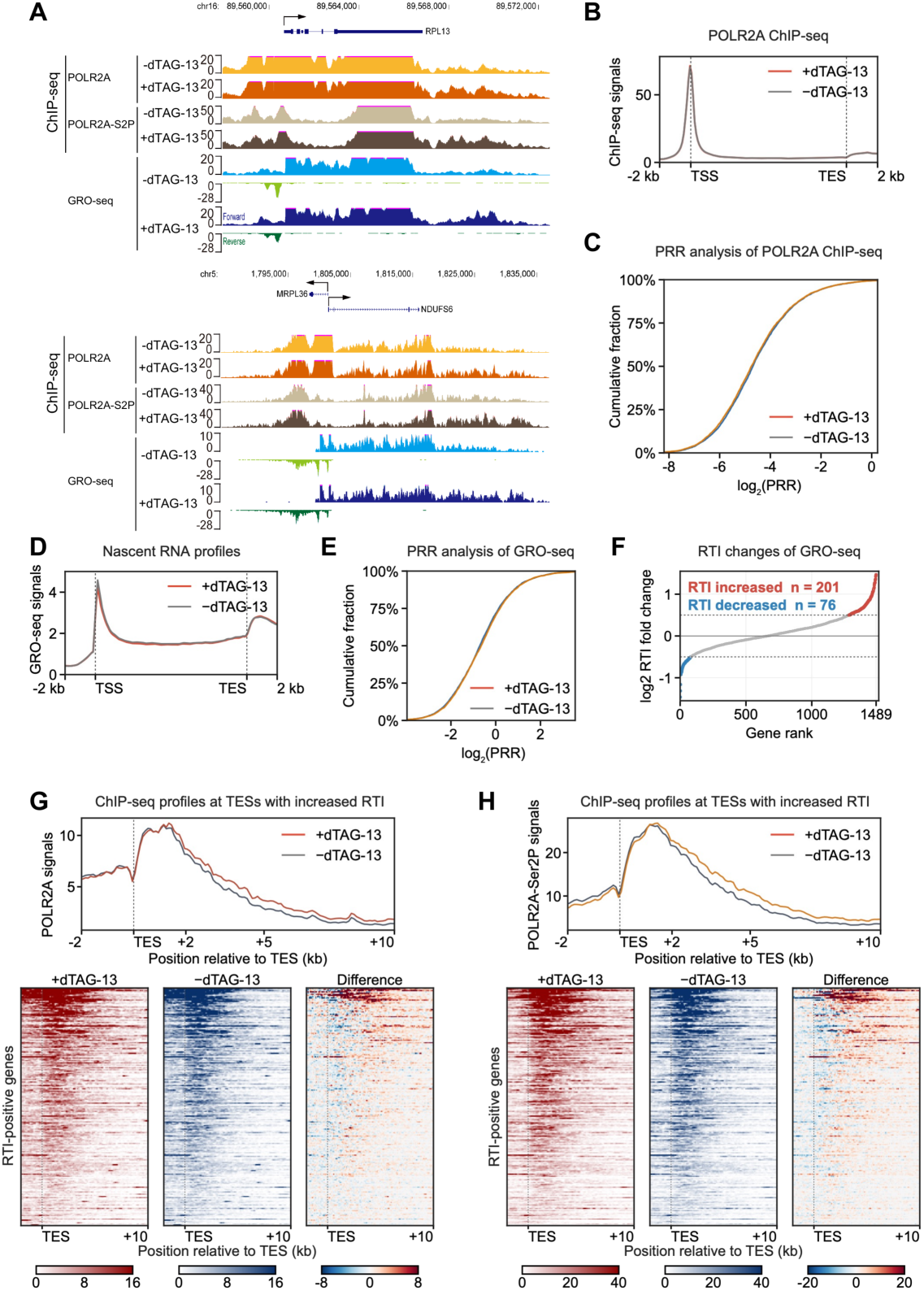
Acute SRSF5 depletion selectively increases downstream transcription without a global pause-release defect. A, POLR2A, POLR2A-S2P and strand-resolved GRO-seq tracks at *RPL13* and the divergent *MRPL36*–*NDUFS6* locus before and after acute SRSF5 depletion. B, Mean POLR2A N-terminal-domain ChIP–seq profile across 4,681 eligible protein-coding genes from 2 kb upstream of the TSS to 2 kb downstream of the TES. C, Empirical cumulative distributions of POLR2A ChIP–seq PRR in the same 4,681 genes. The median change in log_2_(PRR) was −0.059 (0.960-fold), and PRR increased in 40.8% of genes (two-sample Kolmogorov–Smirnov P = 1.51 × 10^−1^; paired Wilcoxon signed-rank P < 2.2 × 10^−16^). D, Mean sense GRO-seq profile across the 4,540 genes that also passed GRO-seq coverage filters. E, Empirical cumulative distributions of GRO-seq PRR in these 4,540 genes. The median change in log_2_(PRR) was 0.033 (1.023-fold), and PRR increased in 52.6% of genes (two-sample Kolmogorov–Smirnov P = 9.81 × 10^−2^; paired Wilcoxon signed-rank P = 1.29 × 10^−5^). F, Ranked mean log_2_ fold change in GRO-seq RTI for 1,489 evaluable genes. Genes with a mean change of at least 0.5 and positive changes in both biological replicates were classified as increased (n = 201); genes with a mean change of at most −0.5 and negative changes in both replicates were classified as decreased (n = 76). G,H, Mean profiles and heat maps of POLR2A (G) and POLR2A-S2P (H) ChIP–seq signal from 2 kb upstream to 10 kb downstream of the TES in the fixed set of 201 GRO-seq RTI-increased genes. The same gene order is used for depleted, control and depleted-minus-control matrices. NC, non-depleted control; KD, acute SRSF5 depletion. Promoter-proximal SRSF5 occupancy among the RTI-increased genes is summarized in Supplementary Table S1.

We next examined whether SRSF5 regulates transcription termination. Interestingly, acute SRSF5 depletion resulted in increased total Pol II occupancy, elongating Pol II signals, and GRO-seq signals downstream of the transcription end sites (TESs) of representative genes, including *RPL13* and *MRPL36-NDUFS6*, indicative of transcriptional readthrough (Figure 6A). To systematically quantify termination defects, we calculated the readthrough index (RTI) genome-wide using GRO-seq signals. This analysis revealed a predominant increase in RTI following SRSF5 depletion, with 201 genes exhibiting increased readthrough compared with 76 genes showing decreased RTI (Figure 6F). Consistently, metagene and heatmap analyses revealed elevated total Pol II and elongating Pol II signals downstream of these affected genes (Figure 6G and 6H).

Among the 201 RTI-increased genes, 51 (25.4%) overlapped a pooled SRSF5 ChIP-seq peak within 2 kb of the selected TSS; this fraction was enriched relative to the remaining genes in the 3,660-gene RTI-eligible background (odds ratio = 1.67, two-sided Fisher’s exact test, P = 0.0039; Supplementary Table S1), supporting a potential association between promoter-proximal SRSF5 occupancy and susceptibility to transcriptional readthrough. Together, these findings demonstrate that SRSF5 selectively facilitates transcription termination at a subset of genes without broadly affecting transcription initiation or elongation. The limited scope of termination defects may reflect functional redundancy among SR proteins or gene-specific requirements for SRSF5-mediated regulation.

## Discussion

Rapid and inducible protein degradation technologies have transformed the ability to interrogate protein function with temporal precision (Ma et al., 2025; Tsai et al., 2024; Zhang et al., 2025). However, our systematic analysis reveals an important limitation of current degron strategies: direct fusion of degron tags to endogenous coding sequences frequently causes unintended reductions in target protein abundance (Figure 1C–I), limiting the generation and interpretation of degron cell lines. This finding is consistent with recent reports describing reduced expression of endogenous degron-tagged proteins in both cultured cells and animal models (Bondeson et al., 2022; Xing et al., 2025; Yenerall et al., 2023), suggesting that tag insertion-associated perturbation represents a general challenge rather than an isolated technical issue. By separating protein production from protein degradation, idTAG overcomes this limitation and provides a complementary strategy for physiological interrogation of endogenous proteins.

idTAG offers two major advantages over conventional degron approaches. First, the Tet-On 3G module enables precise, dose-dependent restoration of protein expression, allowing degron-tagged proteins to be maintained at physiological levels and improving the efficiency of degron cell line generation. Second, independent control of doxycycline-mediated protein synthesis and dTAG-mediated degradation enables flexible tuning of degradation kinetics, ranging from gradual depletion following Dox withdrawal to rapid removal upon ligand-induced degradation. Together, these features establish idTAG as a versatile platform for generating physiologically faithful degron cell lines and achieving precise temporal control of endogenous protein abundance.

Application of idTAG to FUS illustrates the importance of temporal resolution in defining protein function. Previous studies using chronic RNA interference suggested that FUS regulates RNA polymerase II transcription, including effects on Pol II CTD Ser2 phosphorylation (Schwartz et al., 2012). In contrast, acute FUS depletion using idTAG caused minimal changes in global Pol II occupancy, pause release, and nascent transcription (Figure 4). This discrepancy suggests that transcriptional alterations observed after prolonged FUS depletion may arise, at least in part, from secondary consequences of chronic protein loss, including cellular adaptation or stress responses. Our findings therefore refine the current understanding of FUS biology and suggest that, under basal conditions, FUS is not broadly required for transcriptional homeostasis. Instead, FUS may exert more prominent functions in RNA processing, stress granule regulation, and genome maintenance pathways; loss of these FUS functions is strongly implicated in neurodegenerative disease (Birsa et al., 2020; Naumann et al., 2018; Zhao et al., 2026).

Whereas acute FUS depletion had little effect on global transcription, acute SRSF5 depletion revealed a selective role for SRSF5 in transcription termination. Although SR proteins have traditionally been studied as regulators of pre-mRNA splicing, increasing evidence indicates that individual SR proteins can exert distinct transcriptional functions (Long & Caceres, 2008; Zhong et al., 2009). SRSF1 and SRSF2 regulate promoter-proximal pause release through the 7SK-P-TEFb axis (Ji et al., 2013), whereas SRSF5 depletion did not affect pause release but instead caused transcriptional readthrough at a subset of genes (Figure 6). This functional divergence highlights the specialization of SR proteins beyond their shared splicing-related activities. The selective nature of SRSF5-dependent termination defects further suggests that individual SR proteins regulate distinct transcriptional programs rather than serving as globally redundant components of transcription regulation.

Mechanistically, SRSF5 ChIP-seq revealed widespread enrichment of SRSF5 at promoter-proximal regions, with nearly 90% of binding sites localized near transcription start sites. This promoter-associated localization is consistent with previous observations that components of the 3′ end processing machinery, including cleavage stimulation factor and cleavage and polyadenylation specificity factor (CPSF), can be recruited to transcription start sites and remain associated with RNA polymerase II throughout the transcription cycle (Dantonel et al., 1997; Glover-Cutter et al., 2007). Together with recent evidence linking SRSF5 to CPSF7 (Okuda et al., 2025), a core component of the cleavage and polyadenylation machinery, these findings suggest that promoter-bound SRSF5 may contribute to establishing a transcriptional environment that facilitates downstream 3′ end processing and termination.

Several limitations remain. First, although idTAG provides robust control in cultured cells, further engineering will be required to adapt the system for in vivo applications. Second, the molecular basis of SRSF5-mediated transcription termination remains to be directly demonstrated. Third, while acute FUS depletion did not alter basal transcription, potential roles of FUS in stress-responsive or disease-relevant transcriptional programs remain to be investigated. Nevertheless, idTAG provides a powerful framework for dissecting immediate protein functions while minimizing confounding effects caused by chronic perturbation. Beyond establishing a new approach for programmable protein control, our study reveals that acute depletion can uncover distinct biological functions of RBPs, demonstrating that temporal precision is essential for understanding protein function in physiological contexts.

## Materials and methods

### Cell culture and small-molecule treatments

HepG2 and HeLa cells were obtained from the American Type Culture Collection and maintained at 37 °C with 5% CO_2_. HepG2 cells were cultured in MEM (Gibco, #41090101) supplemented with 10% fetal bovine serum (AusGeneX, #FBSCN500-S), penicillin– streptomycin (Gibco, #15140122), non-essential amino acids and sodium pyruvate (Gibco, #11360070). HeLa cells were cultured in DMEM (Gibco, #21013024) supplemented with 10% fetal bovine serum and penicillin–streptomycin.

FUS-idTAG HepG2 and SRSF5-idTAG HeLa cells were cultured with 10 and 20 ng/mL doxycycline (Dox), respectively, for 48 h before depletion. Acute depletion was induced by Dox withdrawal and addition of 0.5 μM dTAG-13 for 6 h (Nabet et al., 2020; Nabet et al., 2018). Control cells retained Dox and received no dTAG-13.

FUS depletion kinetics were measured after Dox withdrawal alone (0, 12, 24, 36 and 48 h), addition of 0.5 μM dTAG-13 with Dox maintained (0, 1, 3, 6 and 12 h), or combined Dox withdrawal and dTAG-13 treatment (0, 1, 3, 6 and 12 h). SRSF5 depletion kinetics were assessed using the combined treatment at the same time points.

### Construction of AID2, dTAG and idTAG donor plasmids

AID2 donors contained approximately 800-bp homology arms flanking the target start or stop codon and an mAID2–mEGFP–FLAG tagging cassette with a P2A-linked blasticidin-resistance marker in pUC19 (Yesbolatova et al., 2020). dTAG donors were generated by replacing mAID2 with FKBP12(F36V). idTAG donors contained an SV40 poly(A) signal, a puromycin-resistance cassette, a TRE3GS promoter and an in-frame FLAG–mEGFP– FKBP12(F36V)–HRV–HA module upstream of the target start codon. Donors were assembled with a homologous-recombination kit (Novoprotein, #NR005). Single-guide RNAs were cloned into pX459, and all constructs were verified by Sanger sequencing.

### Generation and validation of degron knock-in cell lines

AID2 parental HepG2 cells were generated by CRISPR–Cas9-mediated integration of CMV-driven OsTIR1(F74G)–mCherry at the AAVS1 locus (Yesbolatova et al., 2020). For target tagging, donor and pX459–sgRNA plasmids were co-transfected with Lipofectamine 2000. Cells were selected with puromycin (1 μg ml^−1^) and blasticidin (8 μg ml^−1^), and GFP-positive cells were single-cell sorted. Clones were screened by genomic PCR, Sanger sequencing and immunoblotting. idTAG cells were generated by co-transfection of the target-specific donor and pX459–sgRNA plasmid. Dox (100 ng/mL) was maintained during recovery and puromycin selection. GFP-positive single-cell clones were screened by junction and wild-type-allele PCR, and homozygous clones were confirmed by immunoblotting.

### Dual-fluorescence reporter assay

HepG2 cells were transiently transfected with CMV-driven EGFP–FUS–P2A–mCherry or FKBP12(F36V)–EGFP–FUS–P2A–mCherry reporters and analysed by flow cytometry without dTAG-13. EGFP signal was normalized to the co-translated mCherry signal in viable single cells.

### RNA extraction and RT–qPCR

Total RNA was extracted with TRIzol, DNase treated and reverse transcribed with the HiScript III First Strand cDNA Synthesis Kit (Vazyme). Quantitative PCR used Hieff qPCR SYBR Green Master Mix (Yeasen). *FUS* expression was normalized to *ACTB* and wild-type cells by the 2^−ΔΔCq^ method (Livak & Schmittgen, 2001). Three independent biological replicates were analysed.

### Immunoblotting and fluorescence microscopy

Cell lysates were separated by SDS–PAGE, transferred to PVDF membranes and analysed by chemiluminescence. Primary antibodies for immunoblotting were anti-FUS (ABclonal, #A21830), anti-TAF15 (ABclonal, #A9383), anti-HNRNPA1 (ABclonal, #A11564), anti-XRCC6 (ABclonal, #A7330), anti-NONO (ABclonal, #A3800), anti-HNRNPK (ABclonal, #A0772), anti-FLAG M2 (Sigma-Aldrich, #F1804), anti-HA (Cell Signaling Technology, #2999S), anti-SRSF5 (Bethyl Laboratories, #A303-723A) and anti-ACTB (ABclonal, #AC026); ACTB served as the loading control. Band intensities were quantified in ImageJ and normalized to ACTB. Bright-field and mCherry fluorescence images were acquired for the OsTIR1(F74G) parental line.

### Chromatin immunoprecipitation and sequencing

Approximately 1–2 × 10⁷ cells were fixed with 1% formaldehyde for 20 min and quenched with glycine. Chromatin was sonicated to approximately 150 bp and immunoprecipitated with anti-Rpb1 N-terminal domain (clone D8L4Y; Cell Signaling Technology, #14958), anti-Ser2-phosphorylated POLR2A (Abcam, #ab5095) or anti-FLAG M2 (Sigma-Aldrich, #F1804) coupled to Protein A/G magnetic beads.

Immunoprecipitated DNA was reverse-crosslinked, purified and used to prepare VAHTS libraries (Vazyme) for Illumina NovaSeq 6000 sequencing. Two independent biological replicates were analysed per condition.

### GRO-seq

Nuclear run-on reactions were performed for 5 min at 30 °C with ATP, GTP, CTP and Br-UTP (Core et al., 2008). Nascent RNA was extracted, fragmented to approximately 100–200 nucleotides, DNase treated and enriched with anti-BrdU/BrUTP monoclonal antibody-conjugated agarose (Santa Cruz Biotechnology, #sc-32323 AC). Strand-specific libraries were prepared with the NEBNext Ultra II Directional RNA Library Prep Kit (NEB, #E7760L) and sequenced on an Illumina NovaSeq 6000. Two independent biological replicates were generated per condition.

### ChIP–seq processing and peak annotation

All ChIP–seq datasets were processed using the same workflow. Reads were trimmed with Trim Galore v0.6.10, aligned to hg38 with Bowtie2 v2.4.5 and deduplicated with Picard v3.0.0 (Langmead & Salzberg, 2012; Martin, 2011). RPGC-normalized bigWig tracks were generated with deepTools v3.5.4 and used for browser tracks, profiles, heat maps and quantitative analyses (Ramírez et al., 2016).

SRSF5 peaks were called from pooled FLAG ChIP and input replicates with MACS2 v2.2.7.1 at q < 0.05. Blacklist-overlapping peaks were removed, yielding 2,888 peaks, which were annotated with ChIPseeker v1.34.1 using a ±3-kb TSS window (Amemiya et al., 2019; Yu et al., 2015; Zhang et al., 2008).

### GRO-seq processing

GRO-seq reads were trimmed with Trim Galore, depleted of rRNA-derived pairs with Bowtie2 and aligned to hg38 with STAR v2.7.10b using GENCODE v46 annotation (Dobin et al., 2012; Langmead & Salzberg, 2012; Mudge et al., 2024). Uniquely mapped reads were deduplicated with Picard. Strandedness was verified with RSeQC v5.0.1, and strand-specific bigWig tracks were generated with its bam2wig.py utility (Wang et al., 2012). Gene-level counts were obtained with featureCounts v2.0.6 using reverse-strand assignment (Liao et al., 2013).

### Metagene profiles and signal matrices

For metagene and ratio analyses, GENCODE v46 annotation was used (Mudge et al., 2024), and one protein-coding transcript per gene was selected in the order MANE Select, APPRIS principal, Ensembl canonical and longest protein-coding transcript (Morales et al., 2022; Pozo et al., 2022). Unless stated otherwise, transcripts shorter than 2 kb or lacking 2 kb of strand-aware downstream intergenic space were excluded. Condition-level profiles used replicate-averaged bigWig tracks unless stated otherwise. SRSF5 pause-release profiles used 100-bp bins from TSS −2 kb to TES +2 kb, with gene bodies scaled to 10 kb. Profiles for the 201 RTI-increased genes spanned TES −2 kb to TES +10 kb; the same genes and row order were used for POLR2A and POLR2A-S2P.

### Replicate-correlation analysis

Replicate concordance was assessed with deepTools multiBigwigSummary across non-overlapping 1-kb hg38 bins after excluding ENCODE blacklist regions and chrM (Amemiya et al., 2019; Ramírez et al., 2016). GRO-seq forward and reverse signals were combined before analysis. Pearson correlations were calculated from untransformed mean bin signals, and density plots show log1p-transformed signals. Two independent biological replicates were analysed per condition.

### Pause-release and readthrough analyses

For SRSF5, PRR was defined as mean signal from TSS +500 bp to TES −500 bp divided by mean signal from TSS −300 to +100 bp. Protein-coding genes longer than 2 kb, separated from neighbouring genes by >1 kb and containing a control POLR2A promoter peak in both replicates were retained. This yielded 4,681 genes for POLR2A and 4,540 genes for GRO-seq.

SRSF5 GRO-seq RTI was defined as mean strand-resolved signal from 5 to 10 kb downstream of the TES divided by terminal-exon signal. Eligible genes were required to have control GRO-seq TPM > 1, a terminal exon of at least 100 bp and no overlapping gene within 10 kb downstream. Figure 6F included 1,489 genes with control terminal-exon signal ≥1. Within each biological replicate, the RTI change was calculated as the log_2_ ratio of RTI in acutely SRSF5-depleted cells to RTI in non-depleted control cells. Genes were classified as RTI-increased when the mean change across the two replicates was ≥0.5 and the change was positive in both replicates; genes were classified as RTI-decreased when the mean change was ≤−0.5 and the change was negative in both replicates. The 201 RTI-increased genes were used in Figure 6G,H. Promoter-proximal SRSF5 occupancy was defined as overlap of at least one pooled, blacklist-filtered SRSF5 ChIP-seq peak (MACS2 q < 0.05) with the 2-kb interval on either side of the selected TSS. Enrichment among the 201 RTI-increased genes was tested by a two-sided

Fisher’s exact test against the remaining genes in the 3,660-gene RTI-eligible background, comprising genes that met the transcript-selection criteria, had control GRO-seq TPM > 1, had a terminal exon of at least 100 bp and had no overlapping gene within 10 kb downstream of the TES.

### Statistical analysis

Cell-based data are presented as mean ± s.d. from three independent biological replicates unless stated otherwise. Wild-type comparisons used two-tailed unpaired Student’s t-tests, genome-wide distributions used two-sample Kolmogorov–Smirnov tests, and paired per-gene PRR comparisons used two-sided Wilcoxon signed-rank tests. Gene counts and P values are reported in the figures and legends.

## Author contributions

M.K.S., J.K.P., F.Z., and R.X. designed the experiments. M.K.S. performed all the experiments; J.K.P. processed and analyzed all the sequencing data; M.K.S., J.K.P., F.Z., and R.X. wrote the paper.

## Acknowledgements

We thank the staff of the core facility of the Medical Research Institute at Wuhan University for their technical support. This work was supported by the National Natural Science Foundation of China (32371356), the National Natural Science Foundation of Hubei Province (2025AFB648), the Fundamental Research Funds for the Central Universities (2042022dx0003) to R.X., and the Translational Medicine and Interdisciplinary Research Joint Fund of Zhongnan Hospital of Wuhan University (ZNJC202405) to R.X. and F. Z.

---

**Supplementary Table S1.** Promoter-proximal SRSF5 occupancy in the RTI-eligible background.

| Gene group | SRSF5 occupied | Not occupied | Total |
| --- | --- | --- | --- |
| RTI-increased | 51 | 150 | 201 |
| Other RTI-eligible genes | 585 | 2874 | 3459 |
| **Total** | **636** | **3024** | **3660** |

SRSF5 occupancy was defined by overlap of at least one pooled, blacklist-filtered MACS2 peak ( $q < 0.05$ ) with the selected TSS $\pm 2$ kb. The RTI-eligible background comprised 3,660 genes that met the transcript-selection criteria, had control GRO-seq TPM $> 1$ , had a terminal exon of at least 100 bp and had no overlapping gene within 10 kb downstream of the TES. Odds ratio = 1.67; two-sided Fisher's exact test, $P = 0.0039$ .

